## supplemental figures for "Enhancing oscillations in intracranial electrophysiological recordings with data-driven spatial filters"

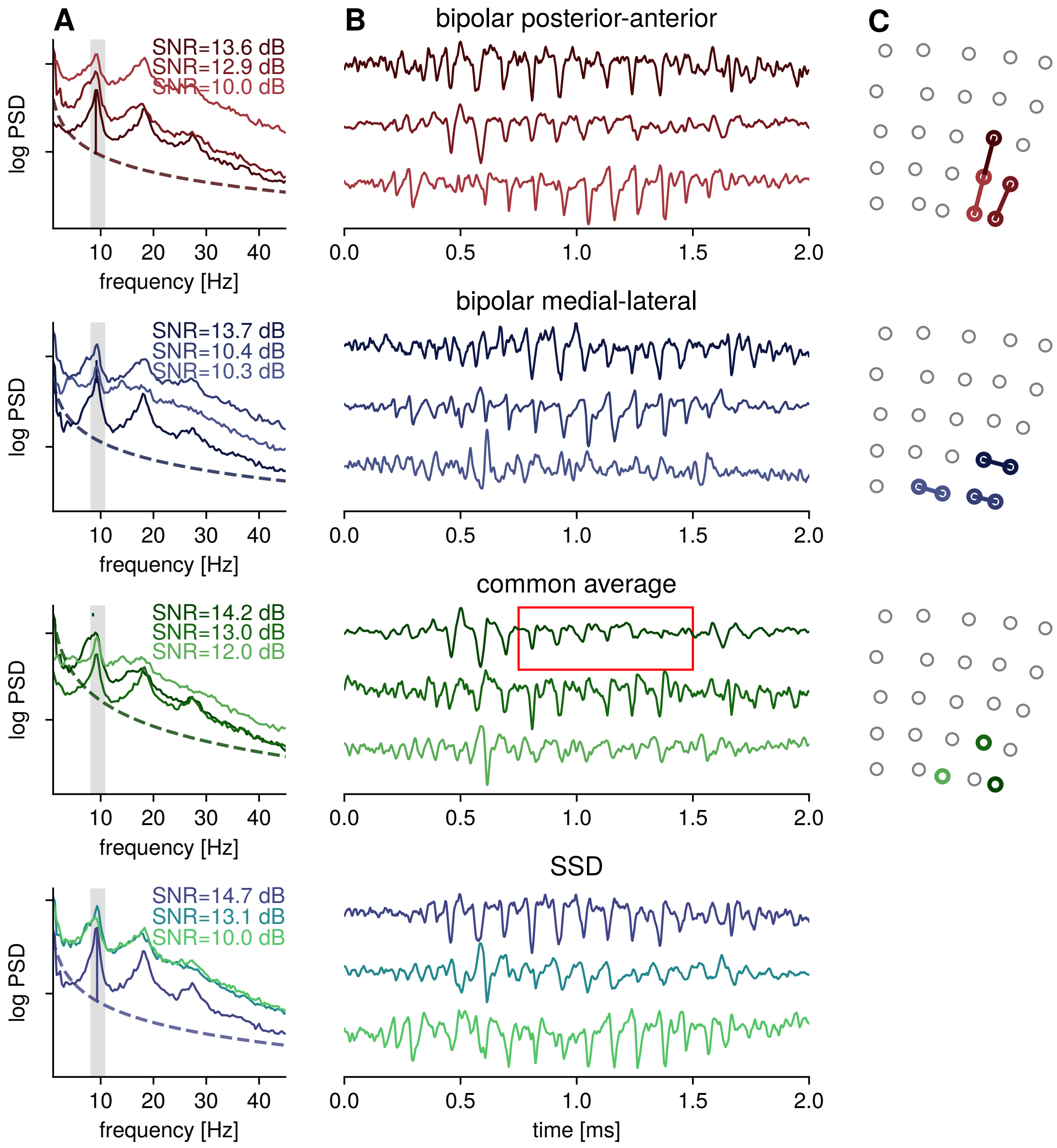


Figure 1: **Example: Four different types of spatial filters.** Shown are respectively for rows, bipolar posterior-anterior referencing, bipolar medial-lateral referencing, common average referencing and SSD data-driven referencing: A) Power spectral densities for three channels with the highest SNR in the highlighted band. The gray bar indicates the frequency band defined as the signal contribution for estimating the SSD spatial filters. The power spectrum shows a spectral peak, with additional harmonic peaks. B) The corresponding signal in the time domain showing oscillatory bursts in the alpha-band, amplitudes are normalized for comparison of time courses. For common-average referencing, the red box marks a time period in which less pronounced oscillations can be seen in the common average reference signals, but the oscillatory power of the constituent SSD components is not decreased. C) Electrode grid showing the origin of the electrode signals, for bipolar signals both reference electrodes are shown, for the common average reference row the center electrodes are shown. For SSD spatial patterns and spatial filters see Fig. 2.

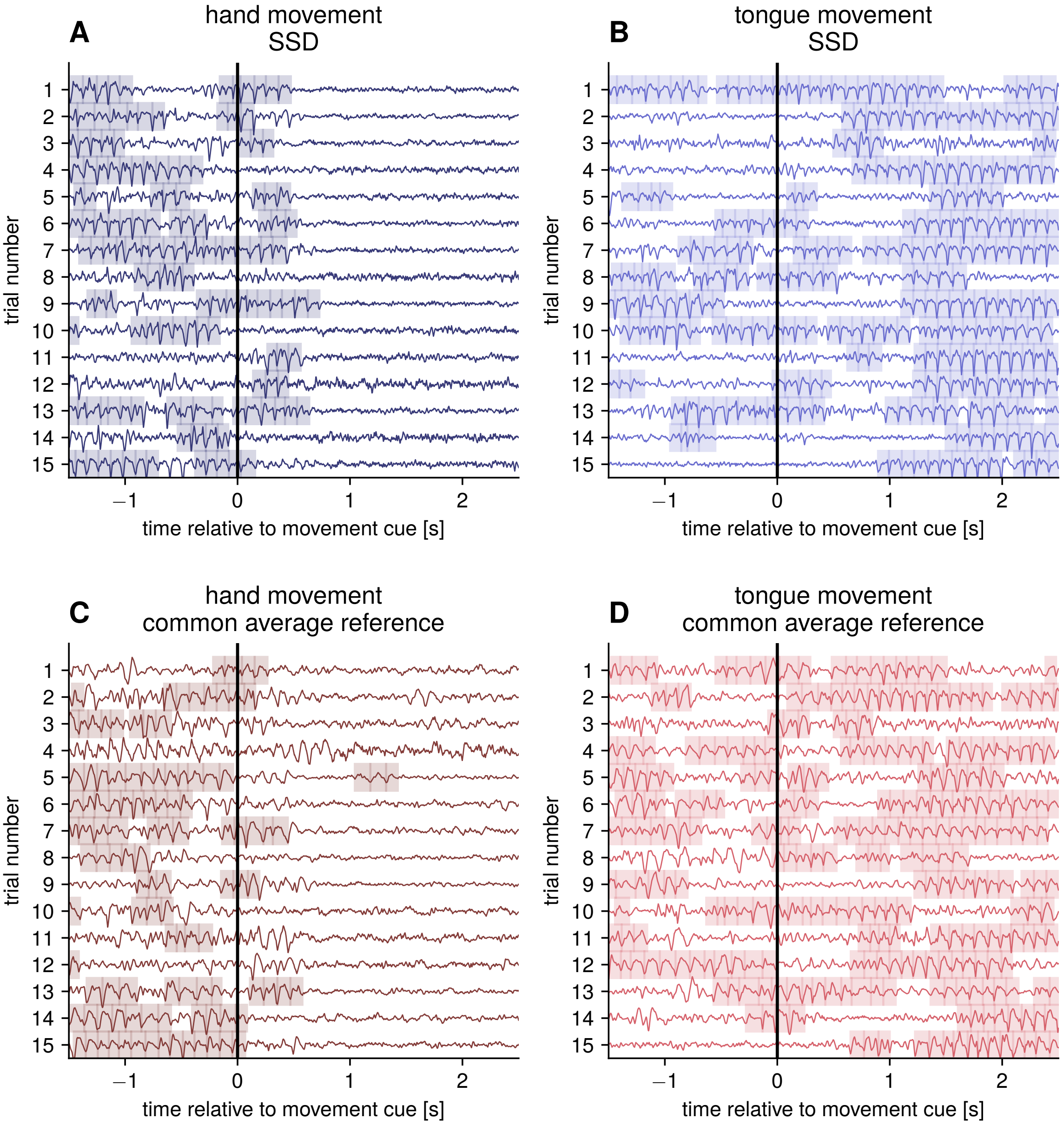


Figure 2: **SSD spatial filters preserve task-related dynamics.** A) Hand-movement trials for the SSD component with highest SNR, with detected burst in the mu-rhythm frequency range highlighted. The event-related desynchronization in the post-cue period is high for hand movements, no desynchronization can be seen for tongue movements, as seen in B). C) The same dynamic can be seen for common average referenced signal for hand movement trials, and tongue movment trials in D). The same burst detection criteria were applied for both referencing methods.
